# Functional and Neurotransmitter Organization of Cortical Gray Matter Remodeling During Pregnancy

**DOI:** 10.64898/2026.11.24.768907

**Authors:** Caroline Hayes, Noemí Rubau-Apa, Tien T. Tong, Laura Pritschet, Hannah Grotzinger, Susana Carmona, Sara L. Kornfield, Rachel Kishton, Magdalena Martínez-García, Golia Shafiei, Emily G. Jacobs, Sheila Shanmugan

## Abstract

Pregnancy is a period of profound neuroplasticity, marked by widespread, spatially heterogeneous reductions in cortical gray matter volume (GMV). Yet, whether this remodeling affects cognition is unclear: subjective memory complaints are common during pregnancy, but objective findings are mixed. Using densely sampled longitudinal magnetic resonance imaging (MRI) across pregnancy, we examined whether gestational GMV change corresponds spatially with normative maps of cognitive function and neurotransmitter receptors. We analyzed 19 T1-weighted MRI sessions of 1 healthy primiparous woman spanning preconception to 36 weeks gestational age. The GMV change map was annotated using 10 cognitive domain maps comprised of 115 Neurosynth terms and 16 positron emission tomography receptor and transporter maps from healthy adults, using dominance analysis and partial least squares (PLS) analysis with spin tests to account for spatial autocorrelation. Total cortical GMV declined by 4.24%. The 10 cognitive domains accounted for significant spatial variation in GMV change (R^2^adj = 0.16; spin *P* = .03), with learning and memory (37.7%) and action (27.3%) accounting for the largest shares of explained variance. PLS revealed a shared spatial axis in which learning and memory were associated with smaller GMV reductions while action was associated with greater reductions (spin *P* = .046). The receptor and transporter maps also accounted for significant spatial variation (R^2^adj = 0.22; spin *P* = .02). PLS identified a significant neurochemical pattern (spin *P* = .009), with greater GMV reductions in regions rich in cholinergic (VAChT) and noradrenergic (NET) markers and smaller reductions in regions rich in serotonergic, GABA_A_, and muscarinic M1 receptors. This receptor-derived pattern was positively associated with social function and negatively associated with action. Together, these findings provide new evidence that pregnancy-related GMV remodeling is organized along functional and neurotransmitter receptor axes, suggesting that maternal brain plasticity is a targeted process that may support selective cognitive adaptation.

## Introduction

Pregnancy is a period of profound neuroplasticity, marked by widespread, spatially heterogeneous reductions in cortical gray matter volume (GMV).^1,2^ These reductions have been associated with maternal attachment,^2^ suggesting an adaptive process that supports the transition to motherhood. Yet, whether this remodeling affects cognition is unclear: subjective memory complaints are common during pregnancy, but objective findings range from subtle deficits that remain within the normative range^3^ to enhanced memory for maternally relevant information.^4^ Whether pregnancy-related GMV remodeling aligns with cortical systems supporting specific cognitive domains may help address this question. Neurotransmitter receptor topography could link these dimensions, as pregnancy hormones act in part through neurotransmitter systems,^5^ and receptor architecture tracks with cortical structure and cognitive specialization.^6^ Using densely sampled longitudinal magnetic resonance imaging (MRI) across pregnancy, we examined whether gestational GMV change corresponds spatially with normative maps of cognitive function and neurotransmitter receptors.

## Methods

This report used the Maternal Brain Project (MBP) dataset^1^ of 19 T1-weighted, 3-Tesla MRI sessions of 1 healthy primiparous woman spanning preconception to 36 weeks gestational age. T1 images were preprocessed using Advanced Normalization Tools and parcellated to the Schaefer 400 atlas. Generalized additive mixed models were used to evaluate GMV change at each parcel across gestation.

To test whether gestational changes in GMV followed functional and neurochemical organization, the GMV change map (Fig 1A) was annotated using 10 predefined cognitive domain maps comprised of a total of 115 Neurosynth terms^7,8^ and 16 positron emission tomography receptor and transporter maps from healthy adults.^6^ For each map set (Neurosynth or neurotransmitter), multiple regression estimated how much of the variance in GMV change the set jointly explained, while dominance analysis quantified individual regressor contributions.^6^ Dominance analysis does not, however, indicate the direction of each regressor’s association with GMV change. As such, we next conducted partial least squares (PLS) analysis to identify multivariate associations between GMV change and receptor topography or functional systems; reliability of PLS loadings was assessed with 95% CI estimated using bootstrap resampling. Throughout, significance was tested using spin tests^9^ with 5,000 rotations to account for spatial autocorrelation, followed by false discovery rate (FDR) correction for multiple comparisons where applicable.

**Figure 1.**
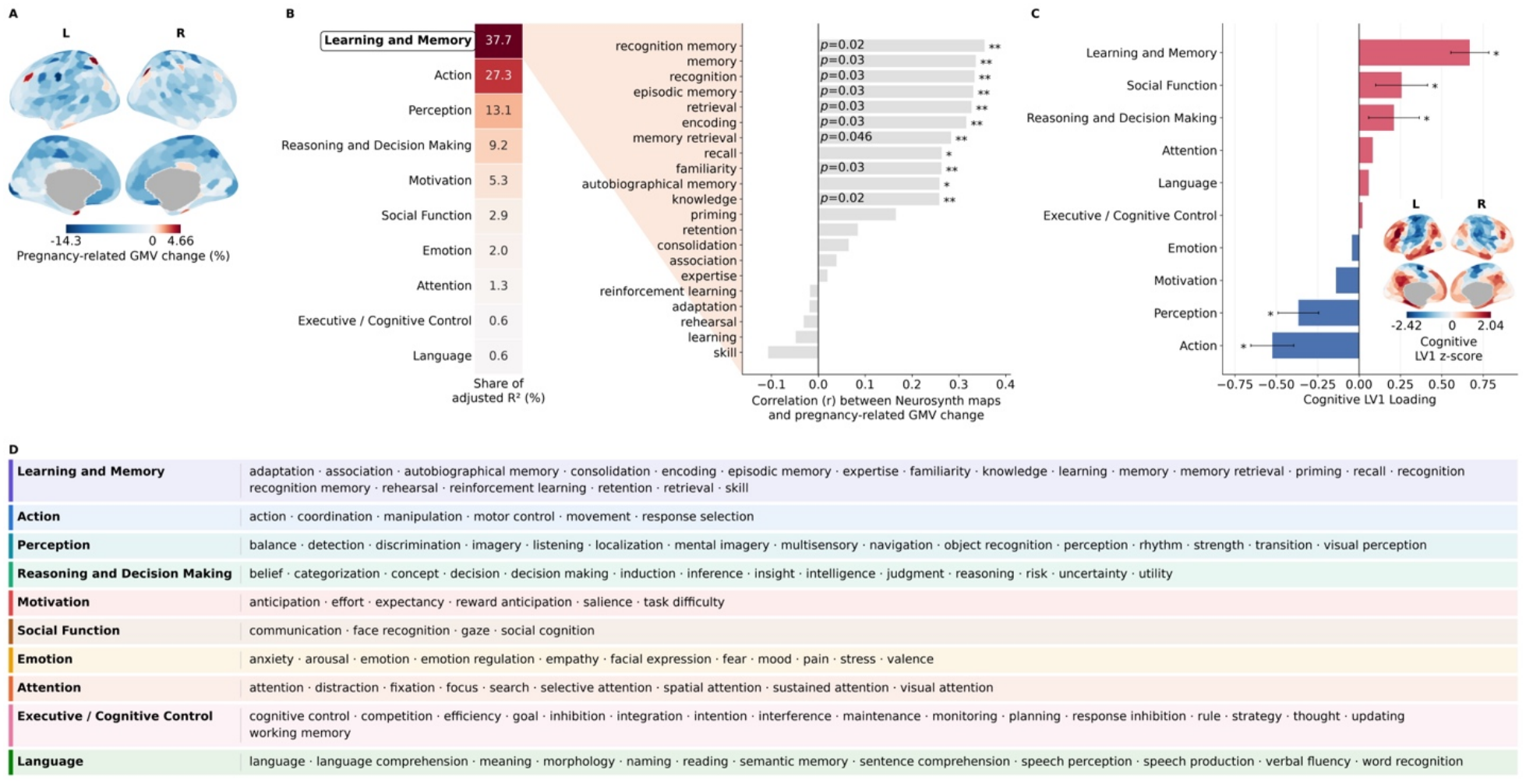
Pregnancy-related cortical gray matter remodeling aligns with a functional axis. To determine whether pregnancy-related gray matter volume (GMV) changes align with the spatial organization of cognitive function, we compared regional GMV changes with Neurosynth-derived meta-analytic activation maps. These maps represented 115 cognitive terms selected based on prior studies and assigned to the 10 domains. **(A)** Generalized additive models fit at each parcel were used to estimate GMV change from preconception to 36 weeks gestation using 19 longitudinal MRI sessions from one primiparous woman. Cortical surface maps depict regional GMV changes (% change at each parcel). Global GMV reductions were observed and were particularly prominent in somatomotor, ventral attention, and dorsal attention regions. **(B)** To identify cognitive domains associated with regional GMV change, we used a dominance analysis to quantify the contribution of each domain to explain spatial variance. Together, the 10 cognitive domains explained significant variation in GMV change (multiple regression adjusted R^2^= 0.16, spin *P* = .03), with learning and memory (37.7%) and action (27.3%) contributing the largest shares of explained variance (left). To further characterize the top-ranked domain, we correlated GMV change with the activation map of each of its 21 constituent terms (right). The strongest positive associations were observed for terms related to memory, recognition, encoding, and retrieval, indicating smaller GMV reductions in regions associated with these processes. Bars represent parcel-wise Pearson correlation coefficients. Statistical significance was assessed using spin-based permutation testing with 5,000 rotations to account for spatial autocorrelation (*spin *P* < .05; **FDR-corrected spin *P* < .05). **(C)** To determine whether GMV remodeling and cognitive functions are organized along a shared spatial axis, we used partial least squares (PLS) regression to identify multivariate patterns of spatial covariance between regional GMV change and the 10 cognitive domains. PLS identified a significant latent variable (LV1; spin *P* < .05), defining a functional axis that differentiated learning and memory, social function, and reasoning and decision-making from action and perception. Inset cortical maps show the spatial expression of this shared GMV–cognitive axis. Positive loadings for learning and memory, social function, and reasoning and decision-making were associated with smaller GMV reductions, whereas negative loadings for action and perception were associated with greater reductions. Error bars denote the 95% confidence intervals for reliable loadings estimated using 5,000 bootstrap resamples. (**D**) Assignment of the 115 Neurosynth terms to the 10 cognitive domains.

## Results

Total cortical GMV declined by 4.24% from preconception to gestational week 36 (R^2^adj = 0.83; *P* < .001). Reductions varied across the cortex and were prominent in somatomotor, ventral attention, and dorsal attention networks (Figure 1A). To determine whether this spatial pattern corresponded to the topography of cognitive functions, we compared regional GMV changes with Neurosynth-derived meta-analytic activation maps representing 10 cognitive domains. Together, these domains accounted for significant spatial variation in GMV change (R^2^adj = 0.16; spin *P* = .03; Figure 1B). Learning and memory (37.7%) followed by action (27.3%) accounted for the largest shares of explained variance.

To explore the processes underlying the largest contributor—learning and memory—we evaluated which specific terms that comprise this domain correlated with the GMV-change map. Gestational GMV change associations with learning and memory were primarily driven by terms related to memory encoding, retrieval, and recognition, as well as broader terms like memory and knowledge (FDR-corrected spin *P* < .05; Figure 1B). PLS analysis of the GMV change map on all 10 cognitive domains further identified a significant latent variable (LV1; spin *P* = .046; Figure 1C), revealing a shared spatial axis linking GMV remodeling to cognitive function. In this axis, learning and memory were associated with smaller GMV reductions while action was associated with greater reductions.

We next examined whether gestational GMV changes corresponded to neurotransmitter receptor distributions. Together, the 16 receptor and transporter maps accounted for significant spatial variation in GMV change (R^2^adj = 0.22; spin *P* = .02; Figure 2A), with the largest contributions from vesicular acetylcholine transporter (VAChT; 33.3%) and serotonin 5-HT1A receptors (18.0%). PLS identified a significant neurochemical pattern (spin *P* = .009; Figure 2B), with greater GMV reductions associated with cholinergic markers (VAChT) and noradrenergic markers (norepinephrine transporter [NET]) and smaller GMV reductions associated with regions with higher density of serotonergic receptors, GABA_A_ receptors, and the muscarinic cholinergic M1 receptor.

**Figure 2.**
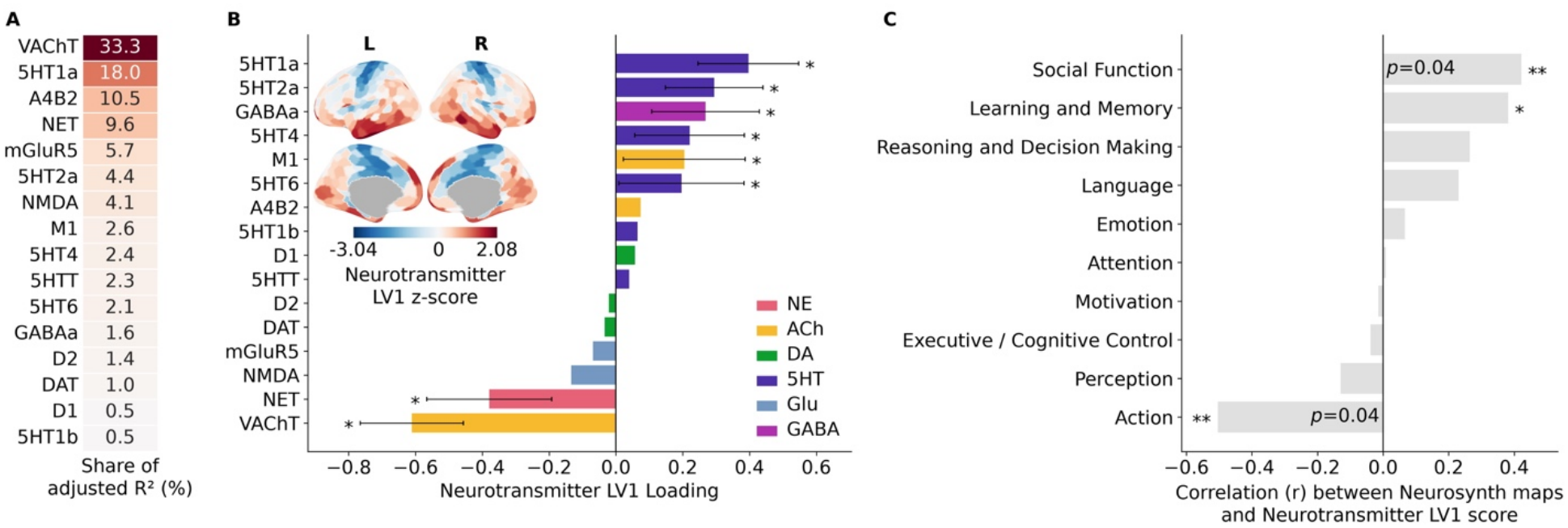
Pregnancy-related cortical gray matter remodeling aligns with neurotransmitter receptor organization. To determine whether pregnancy-related gray matter volume (GMV) change is aligned with the spatial organization of neurotransmitter systems, we compared the regional GMV changes seen in Fig 1A with 16 normative positron emission tomography-derived receptor and transporter maps spanning six neurotransmitter systems. **(A)** Dominance analysis quantified the relative contribution of each receptor or transporter map in explaining regional variation in GMV change. Together, the 16 maps explained significant variation in GMV change (adjusted R^2^= 0.22, spin *P* = .02), with vesicular acetylcholine transporter (VAChT) and serotonin 5-HT1A receptors contributing the largest shares of explained variance. **(B)** To determine whether GMV remodeling and neurotransmitter receptor distributions are organized along a shared spatial axis, we used partial least squares (PLS) regression to identify multivariate patterns of spatial covariance between regional GMV change and the 16 receptor and transporter maps. PLS identified a significant latent variable (LV1; spin *P* = .009), defining a neurochemical axis that differentiated cholinergic and noradrenergic markers from serotonergic, muscarinic, and GABAergic signatures. Inset cortical maps show the spatial expression of this shared GMV-receptor axis. Bars depict receptor loadings, reflecting the strength and direction of each receptor’s association with the latent axis. Negative loadings for VAChT and NET were associated with greater GMV reductions, whereas positive loadings for serotonergic, GABA_A_, and M1 receptors were associated with smaller reductions. Error bars denote the 95% confidence intervals for reliable loadings estimated using 5,000 bootstrap resamples. **(C)** To determine whether the neurochemical pattern followed the functional organization of cognitive functions, we correlated receptor-derived LV1 scores with Neurosynth activation maps representing 10 cognitive domains. LV1 was positively associated with social function and learning and memory and negatively associated with action. Bars represent parcel-wise Pearson correlation coefficients. Statistical significance was assessed using spin-based permutation testing with 5,000 rotations to account for spatial autocorrelation (*spin *P* < .05; **FDR-corrected spin *P* < .05).

To characterize the functional profile of this receptor-derived pattern, we compared the receptor latent-variable map with the 10 cognitive domain maps using Pearson correlations. The receptor-derived pattern showed a significant positive spatial association with social function and a significant negative association with action (FDR-corrected spin *P* < .05; Figure 2C), such that regions linked to social function showed smaller GMV reductions and regions linked to action showed greater reductions.

## Discussion

In this densely-sampled, longitudinal study, pregnancy-related GMV reductions were organized along functional and neurotransmitter axes. More prominent GMV reductions were observed in action-related cortex and regions rich in cholinergic and noradrenergic markers, while lesser reductions were seen in memory-related cortex and regions rich in serotonergic, GABA_A_, and M1 receptors.

Notably, although acetylcholine is often linked to memory,^10^ regions with high VAChT density showed greater GMV reductions while memory-related regions showed relatively smaller GMV reductions. Given the roles of cholinergic and noradrenergic systems in motor learning and action execution,^11^ remodeling in regions associated with these systems may reflect adaptation to evolving sensorimotor demands of changes in body mass, weight distribution, posture, and balance during pregnancy.

Although subjective cognitive complaints are common during pregnancy, objective evidence of cognitive change is modest,^3^ and our findings suggest that memory-associated cortex may be relatively preserved in some pregnancies. Exploring inter-individual variability in perinatal brain remodeling is thus warranted, as other studies have reported prominent remodeling in association regions. Future studies with larger samples that incorporate cognitive assessments are needed to determine whether these spatial patterns are reproducible and how they relate to subjective and objective cognitive changes during pregnancy. Together, these findings provide new evidence that pregnancy-related GMV remodeling is organized along functional and neurotransmitter receptor axes, suggesting that maternal brain plasticity is a targeted process that may support selective cognitive adaptation.

